# Distinct clonal dynamics and interactions within the microenvironment near tumor stroma interfaces in rare histologic variants of bladder cancer

**DOI:** 10.64898/2026.03.21.713423

**Authors:** Luisa Quezada Ziesse, Sophia Bhalla, Antara Biswas, Vignesh Pakiam, Greg Riedlinger, Saum Ghodoussipour, Subhajyoti De

**Affiliations:** Rutgers Cancer Institute, Rutgers the State University of New Jersey, New Brunswick, NJ 08901, USA

## Abstract

Rare histologic variants of bladder carcinoma, such as squamous cell and neuroendocrine carcinoma, generally have a worse prognosis compared to pure urothelial carcinoma (PUC), but the underlying molecular determinants are not well understood. We developed a novel computational genomics framework to characterize the dynamics of tumor-stroma-immune interactions at tumor borders and interiors from spatial transcriptomics data. We profiled bladder carcinoma samples of different histological variants on the Visium platform. Differences in clonal phylogeny and spatial heterogeneity of major subclones between the samples suggested disparate clonal spatiotemporal dynamics and interaction with stromal and immune compartments - which was notably prominent at the tumor-stroma interface. Our framework captured immune heterogeneity in the tumor microenvironment, including variations in the presence and architecture of tertiary lymphoid structures. Our analyses further indicated that there are substantial histology-specific differences in cell type composition, clonal spatial heterogeneity, inter-cellular signaling, and cellular processes. These variations collectively suggest divergent mechanisms of microenvironment remodeling across bladder cancer histologies. Cell-free DNA profiling from liquid biopsy captured tumor and microenvironment signatures from tumor boundaries and interiors, potentially allowing for tracking clonal dominance non-invasively. Our method tracks the trajectory of neoplastic disease in bladder cancer samples while identifying aggressive features.

## 1. Introduction

Bladder cancer (BC) affects over 1.6 million people worldwide with >0.5 million new cases and >0.2 million deaths each year. [1] While most patients present with localized disease that is curable, the heterogeneity of tumor biology presents challenges in risk stratification and risk adapted treatment recommendations. Pure urothelial carcinoma (PUC) is the most common histology, with about 90-95% of cancers originating from the transitional cells that line the bladder, followed by non-urothelial histologies, such as squamous cell carcinomas (SCC) (2-5%) which develop from squamously differentiated transitional cells, adenocarcinomas and sarcomas [2]. Moreover, PUC can differentiate into variant histologies in 25% of the cases, which are typically associated with worse prognosis [3]. While most research has focused on the study of PUC, which has established treatment guidelines, the remaining 5% of rare histological types remain widely understudied. Even in PUC, a limited understanding of intratumoral heterogeneity can influence oncologic outcomes. The high heterogeneity across samples and the rarity of variant histologies presents unique challenges in pathological classification, clinical presentation, risk stratification, and by extension, therapeutic decision making.

Previous studies have aimed to characterize differences between conventional PUCs and histological variants using transcriptomic approaches, [4, 5] but they have mainly focused on functional analyses and have failed to explore the dynamics within the tumor microenvironment (TME). Cancer cells don’t function in isolation and are greatly influenced by other cells present in the TME. Immune cells, cancer associated fibroblasts (CAFs) and endothelial cells are some cell types whose interaction influences cancer progression. Recent cancer therapies have found success in targeting cells in the TME [6, 7] showing alternative ways to combat the disease and highlighting their role in tumor growth. Because of this, there is a need to study not only the cancer cells but also their interaction with the tumor stroma.

Identifying regions with high aggressive activity within the tumor-non-tumor interface can give insight into what pathways are important for tumor progression and potentially identify targets for a specialized mode of care. Spatial transcriptomics (SpaT) has emerged as a powerful tool in cancer research, allowing for the visualization of gene expression patterns within the spatial context of tumor tissues, and enabling the understanding of cellular dynamics of the TME. Although SpaT lacks single cell resolution, knowing the interaction occurring spatially within a tumor can greatly increase our understanding of the disease biology. Independent studies have identified tumor interface and cell signaling inside a tissue sample respectively [8, 9]. However, methods to detect active tumor-stroma borders and the cellular signaling occurring within them remain to be systematically developed.

To address these gaps, we developed CALISTA, an acronym for **C**lassification of **A**ctivity of **L**ocal **I**nterface of **S**troma and **T**umor-signatures using Spatial Transcriptomics **A**nalysis to identify highly active regions within the tumor-stroma interface. Additionally, we established a workflow to analyze clonal dynamics and its interactions with the TME across the interface. Here we applied the framework to analyze the transcriptomic profile of rare bladder cancer histological types by performing SpaT on tumor biopsies. By first identifying the active tumor border, we were able to better characterize the specific interactions that occur at the border and how these can shape the overall trajectory of the disease in different types of both pure and common BC tumors.

## 2. Results

### 2.0 CALISTA Overview

We develop CALISTA as a resource to identify local tumor-immune-stroma interfaces within the tumor microenvironment and uncover tumor clonal dynamics, as well as the functional and spatial heterogeneity in cancer hallmarks and anti-tumor response at tumor-stroma boundaries and interior using spatial transcriptomic data from the 10X Visium platform. We argue that dynamic tumor-immune-stromal interactions at the active, passive, and regressive tumor-stroma boundaries involving multiple different cell types, reflect emergent properties associated with fundamental cancer hallmarks, that have relevance for predicting overall aggressiveness of the tumor.

CALISTA has six main modules (**Figure 1**). Module-A consists of multiple functions to define the stroma-tumor local border by analyzing the spatial patterns of malignancy-associated signatures within a sample. Module-B classifies the border into active, passive or intermediate zones. Active zones could be further classified as progressive and regressive, utilizing cancer hallmark signatures. The remaining modules employ existing strategies to analyze the interactions and dynamics occurring at the identified tumor-stroma interfaces. Module-C makes inferences on major subclones and their localization relative to the active and passive zones and interior. Module-D identifies major stromal cell types, their spatial heterogeneity, and localization relative to the tumor margins and interior. Module-E maps inter-cellular signaling and interactions involving subclones. Module-F projects the likely trajectory of cell populations of interest in pseudotime at the tumor-stroma margins, reflecting the tumor dynamics.

**Figure 1.**
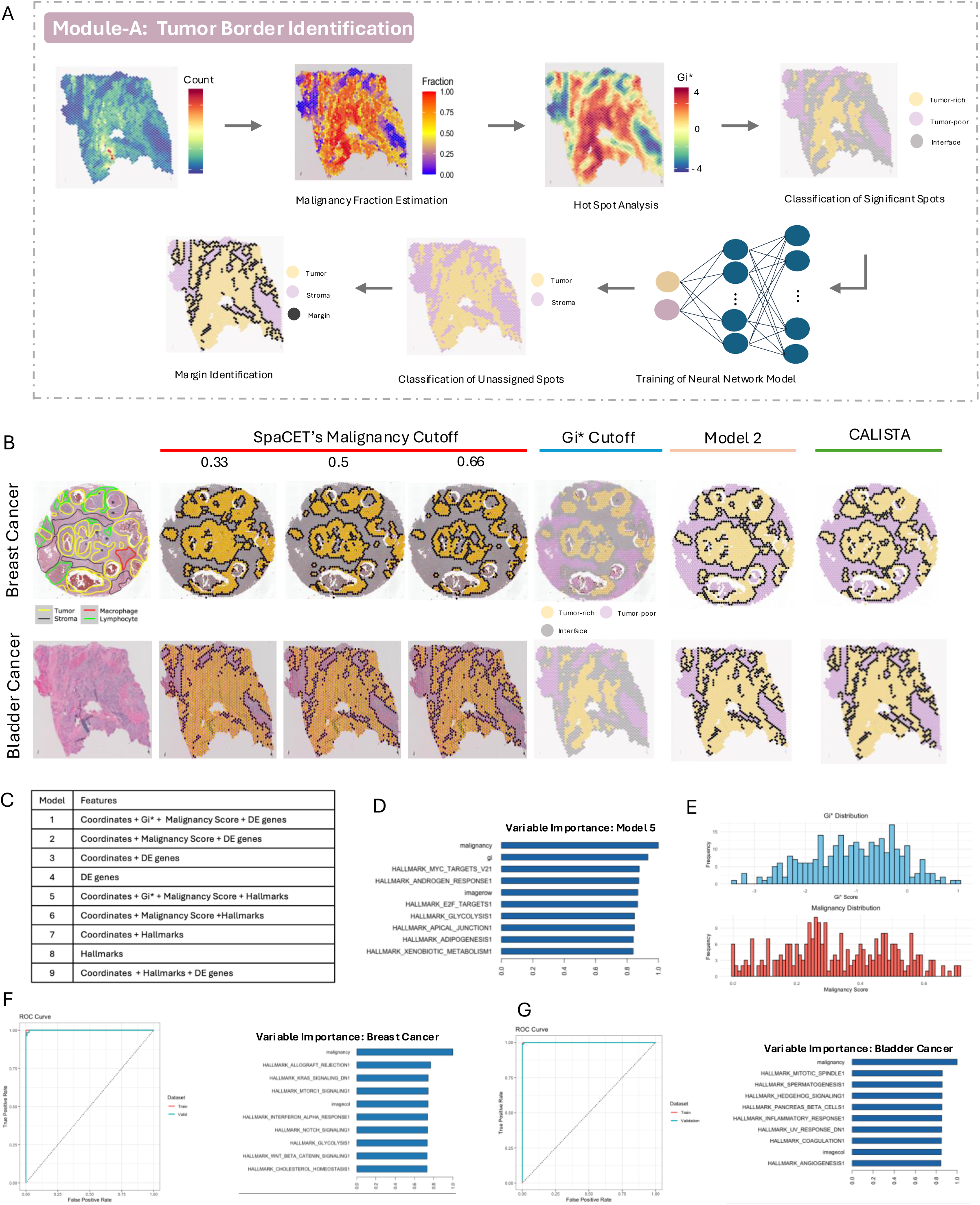
A) Steps for the identification of local tumor borders. B) Comparison of different methods for tumor - nontumor borders across representative in house bladder cancer sample and out house breast cancer sample. C) Table of different models trained for classification of interface spots. D) Important Variables extracted from model 1, bladder cancer sample. E) Distribution of malignancy score and Gi* score across the identified border by model 2 for bladder cancer sample. F) ROC curve and feature importance of model 2 for the bladder cancer sample. G) ROC curve and feature importance of model 2 for the breast cancer sample.

First, we show CALISTA’s capability for active tumor-stroma interface identification through the benchmarking of Module-A and B using in-house and out-house samples. Once we have proven the robustness of our model, we shift our focus to in-house samples obtained from rare and common BC histological subtypes. We show how Modules C through F aids in the understanding of the different subtypes and validates interface classification through the findings within. Finally, we assess these inferences by spatial transcriptomic profiling and liquid biopsy sequencing, as discussed later.

### 2.1 Identification of Local Tumor-stroma interfaces from spatial transcriptomic data

Module-A accepts as input the expression matrix and coordinates provided by the 10X Visium spatial transcriptomics platform. It transforms the spatial transcriptomic data as a Seurat [10, 11] object and extracts multiple tumor biogeographical features that have relevance for tumor stroma boundaries and interiors, which are used in a deep learning-based classifier model to annotate the tumor interfaces and other regions (**Figure 1A**). CALISTA was tested on 9 bladder cancer samples collected inhouse and 3 publicly available SpaT samples from different cancer cohorts. As representative examples, we show the results of Module-A for a bladder cancer sample (S1) and a breast cancer sample (OS1) (**Figure 1 B-G**).

CALISTA calculates a malignancy score to identify cancerous spots, using the approach implemented by SpaCET [8]. The regions with high malignancy scores (close to 1) are tumor rich, while those with low scores (close to 0) are tumor poor. It also computes the Getis-Ord* (*G*i*) [12] statistic, which evaluates the spatial association of a feature based on distance metrics and highlights regions where local clustering occurs. A region with high malignancy score and *G*i* statistic could be considered as tumor-dense, while regions with low scores could be considered as tumor-poor. Local tumor borders may have moderate or low values for these features.

#### 2.1.1. Benchmarking and validation

We evaluated performance of the key CALISTA module for tumor-nontumor border inference by comparing their predictions against pathological annotations using several alternative strategies (**Figure 1B**). As employed by other groups [8], we first determined the tumor-stroma interface by selecting a strict cutoff for the malignancy score, starting with the default of 0.5 and altering it incrementally. Since tumors from different histology have inherently different tumor purity, cutoffs matched histopathological annotations at varying degrees. Even within the same tissue, no single cutoff matched annotations perfectly but rather different cutoffs matched distinct sections of the tumor with higher accuracy.

Through the incremental increase of the malignancy’s cutoff from 0.33 to 0.66 we noticed the predicted tumor border shrinking and expanding. Certain tumor and non-tumor regions remained constant, while areas surrounding the borders changed with each cutoff. We hypothesized that these fluctuating regions exhibited both tumor and non-tumor signals and that malignancy score alone was not enough to classify them into a particular region. As a result, we aimed to identify the non-fluctuating true tumor and non-tumor regions by computing the *G*i* statistics on the malignancy score. This statistic evaluates the spatial association of the feature and infers tumor hotspots and cold spots in a data-driven spatial-context-aware manner.

Based on the spatial distribution and local clustering of the malignancy score, the Gi* score identifies hotspots and cold spots with tumor-rich and tumor-poor regions. This statistic measures both the value of a feature and it’s clustering spatially, as a result statistically significant regions are found in the center section of the tumor-rich and tumor-poor regions. Regions in between these can be thought of as an interface region with both stroma and tumor signals. When comparing the identified interface region to the tumor-border plots determined through the malignancy score, we noticed that fluctuating border spots belong to this region. Our next goal was to use the annotated tumor-rich and tumor-poor regions to train a deep learning-based classifier model that would predict the status of the interface spots on the slide. The boundaries of the tumor and non-tumor regions would be annotated as the local tumor-stroma interfaces.

The classifier model is trained in a tissue dependent manner. Hotspot analysis can classify about half of the tissue. This resulted in around 700-2400 datapoints per sample for model training. The classifier was tested on 9 different inputs, with slightly different features that consisted of a mixture of Malignancy Score, spot coordinates, Cancer Hallmarks Scores and Differentially Expressed (DE) genes between the tumor-rich and tumor-poor regions (**Figure 1C**). Models trained on the Gi* statistic and Malignancy score as input features had perfect classification for both training and validations set (AUC: 1). For these models, variable importance extraction placed both features as top determinants with a relative importance close to 1 (**Figure 1D**). Since our goal was to classify spots whose Gi* statistic scores are not significant, we excluded this feature from the model.

Models 1-4, which included DEs as input features, had generally good performance (AUC: ∼ 0.995), however the runtime was considerable, particularly for samples with more DE’s or spots sequenced. On the other hand, models which used Hallmarks instead of DEs, had a significantly faster runtime and slightly higher accuracy (AUC: ∼0.999). Model 9, which focuses only on the sample’s intrinsic transcriptomic signature by combining DEs and Hallmarks scores but excluding the Malignancy score, had comparable accuracy (AUC: ∼ 0.997) though still long runtime. However, this model helped demonstrate that tumor and stroma regions can be determined from intratumor information alone without the need for correlating signatures to an existing database. Model 6 which consisted of the coordinates, malignancy and Cancer Hallmarks scores between the two groups had the highest overall performance for (i) the outhouse cohort (ii) bladder cancer cohort and (iii) overall good accuracy across both cohorts (AUC > 0.99). When comparing histopathological annotations, we could observe the model matched well. To reduce runtime, increase accuracy, and avoid overfitting Model 6, which consisted of the cancer hallmark scores instead of DEs, was chosen as the classifier model (**Figure 1F-G**).

Classifying spots through this method revealed slightly different tumor-stroma regions than by using arbitrary cutoffs for either malignancy or Gi* statistics. Border spot labeling was assigned to stroma spots with neighboring tumor cells. Identified border regions exhibited a wide range of malignancy scores (S1: 0 - 0.7) and belong to both hot and cold regions (S1: −3 - 1) (**Figure 1E**). The lack of specificity for these two scores highlights the need for a more robust identification method than using a set standard cutoff. According to expert histopathologist observations, using malignancy cutoff alone exhibited accurate results in identifying lone cancer cells inside non-tumor tissue, however it failed to identify continuous borders that cleanly divided tumor and non-tumor tissue. Since our main aim was to identify the interactions within these continuous borders, we decided to use a more complex method for classification.

Expert histopathological annotations were used to identify the tumor-stroma border in H&E-stained images. These annotations were used to validate CALISTA’s identification of the interfaces. There appeared to be high concordance between the expert annotations and CALISTA’s output. When evaluating on samples that lacked histopathological annotations, we noticed that CALISTA identified the topologically distinct regions shown in the sample’s staining. Our method identified tumor-stroma regions using transcriptomic signatures that were able to match histopathological annotations.

CALISTA’s approach provides a rational, data-driven border inference, which proves advantageous over alternative approaches. It avoids border-inference based on user-input, as implemented in SpaCET, and thereby offering scalability and objective annotation. Additionally, feature importance extraction of the classifier model can give insight into pathways driving tumor progression that can be targeted for therapeutic response.

### 2.2 Classification of Active and Passive tumor-stroma interfaces

Next, we used Module-B of CALISTA to characterize the identified local tumor borders by analyzing cellular processes and cancer hallmark signatures to annotate the zones with elevated pro- or anti-tumor activities (**Figure 2A**). Since our main goal was to characterize interactions between the tumor-non-tumor border, only the margin spots with both tumor and non-tumor neighbors were selected for this analysis. Spots are clustered into three groups using k-means clustering based on the activity of the cancer hallmark pathway signatures. Based on the scree plots of the k-mean clustering and biological interpretation we identified 3 dominant clusters indicating High, Medium, and Low cancer activities (**Figure 2B**). Cancer hallmarks such as Epithelial Mesenchymal Transition, Complement, Coagulation, Hypoxia, and Angiogenesis pathway signatures were the dominant features driving the separation among the clusters (**Figure 2C-D**). Cluster-1, indicating high interface activity, had elevated expression of aggressive cancer hallmarks such as, Epithelial Mesenchymal Transition, TGFb signaling and angiogenesis among others. Cluster-2 had intermediate levels of the pathway activities, while cluster-3 with low interface activity, had low expression of these pathways, indicating passive interfaces.

**Figure 2.**
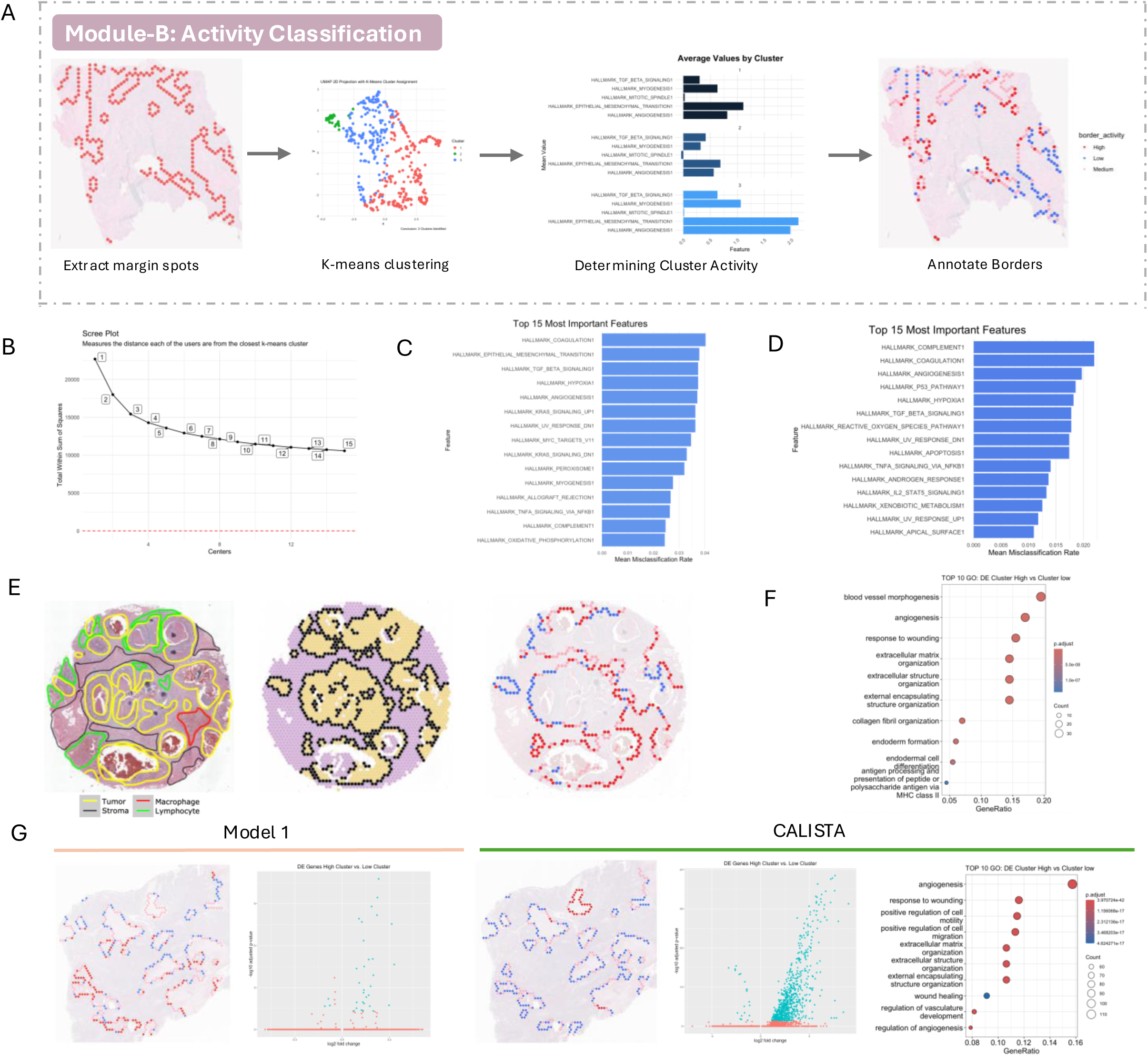
A) Steps for classification of border spots into active, intermediate and passive zones. B) Scree plot from k-means clustering of bladder cancer sample. C) Top 15 most important features extracted from k means clustering of border zones for the representative bladder cancer sample. D) Top 15 most important features extracted from k means clustering of the representative breast cancer sample. E) Expert annotation, predicted tumor - nontumor border, classification of local borders by activity for out-house breast cancer sample. F) Top GO terms of the identified DE’s between active and passive zones in breast cancer sample. G) Classification of tumor-nontumor border zones using 2 different methods and volcano plots of the DE genes between active and passive border zones of in-house bladder cancer sample G.

To verify correct identification of high and low activity regions in the tumor-stroma interfaces, we performed a Differential Expression (DE) Analysis of Active vs Passive clusters (**Figure 1G**). Volcano plots revealed ample amounts of significantly DE genes for almost every sample. Gene ontology (GO) analysis of these genes revealed that most of the genes with altered expression were involved in motility, locomotion, angiogenesis, extra cellular matrix (ECM) remodeling and immune response (**Figure 2F**) - which suggests that our tumor-stroma interface classification can indicate local invasiveness and tumor dynamics. In some cases, the entire tumor-stroma interface was active or passive, while in a few cases there was a gradual transition in the level of activity across an extended margin. Although spatial coordinates were not used during the clustering, local continuity of the interface category suggested that technical noise or other biases in the interface classification is likely low. Downstream analysis revealed how various hallmarks exhibit high autocorrelation with surrounding spots sharing similar scores and slowly fading out, validating the similar behavior observed in contiguous interface sections.

For further validation, we obtained data on the breast cancer sample from Ru et al. [11] whose curated histopathological annotation showed active interfaces surrounding identified macrophages regions and passive interfaces near lymphocyte aggregates (**Figure 2E**). Macrophages often have tumor promoting effects while lymphocytes are typically part of the body’s immune response to fight cancer [13]. Previous studies have identified lymphocyte infiltration in breast tumors to be a prognostic indicator, with high infiltration correlating with overall survival [14]. CALISTA’s ability to classify tumor interfaces surrounding macrophages to be more active while those near lymphocytes to be more passive in breast cancer further highlights the efficiency of our method for identifying active tumor borders without the need for previous annotation of cell types within the sample.

We compared alternate strategies to identify and annotate the interface spot clusters such as malignancy cutoff, Gi* cutoff and different starting input for model training to further assess the robustness and biological relevance of our interface classification. While each method identified similar looking interfaces, the slight differences in composition lead to formation of different clustering which were unable to identify regions with high and low activity as seen by the DE and GO plots. As an example, we show BC sample 6 (**Figure 2G**), whose clustering of interface spots show high sensitivity. The Model 1 fails to identify an interface region whose highly active spots are significantly different to the less active counterparts. These analyses highlight the robustness of our default method and underscore the need to establish well defined interfaces before identifying the active regions.

The modules C-F of CALISTA further reveal the tumor clonal dynamics, and cell-cell signaling at or near the tumor stroma interfaces, as well as those away from the interfaces. We apply CALISTA to a cohort of BC samples from different histological subgroups, as discussed below, that reveal complex and synergistic pro- and anti-tumor activities in the tumor local margins and core regions, and tumor subtype-specific differences therein. These analyses further support the robustness of the interface identification and classification strategies outlined above, and their significance in the context of tumor dynamics.

### 2.2 Clinical and histopathological heterogeneity in the BC cohort

To examine the heterogeneity exhibited across BC tumors, we collected samples belonging to PUC and rare histologic variants (**Figure 3A**). We performed spatial transcriptomics profiling of 5 rare bladder tumor specimens of different histologic subtypes and their corresponding NATs, and analyzed their transcriptomics landscapes together with 10X spatial transcriptomic data from 3 PUC samples and one SCC sample from Biswas et al. The rare tumor specimens included one high-grade small cell neuroendocrine carcinoma (SCNC), 2 primitive neuroectodermal tumor (PNET) samples collected from the same patient, and 2 invasive keratinizing squamous cell carcinoma (KSCC) samples from bladder cancer patients.

**Figure 3.**
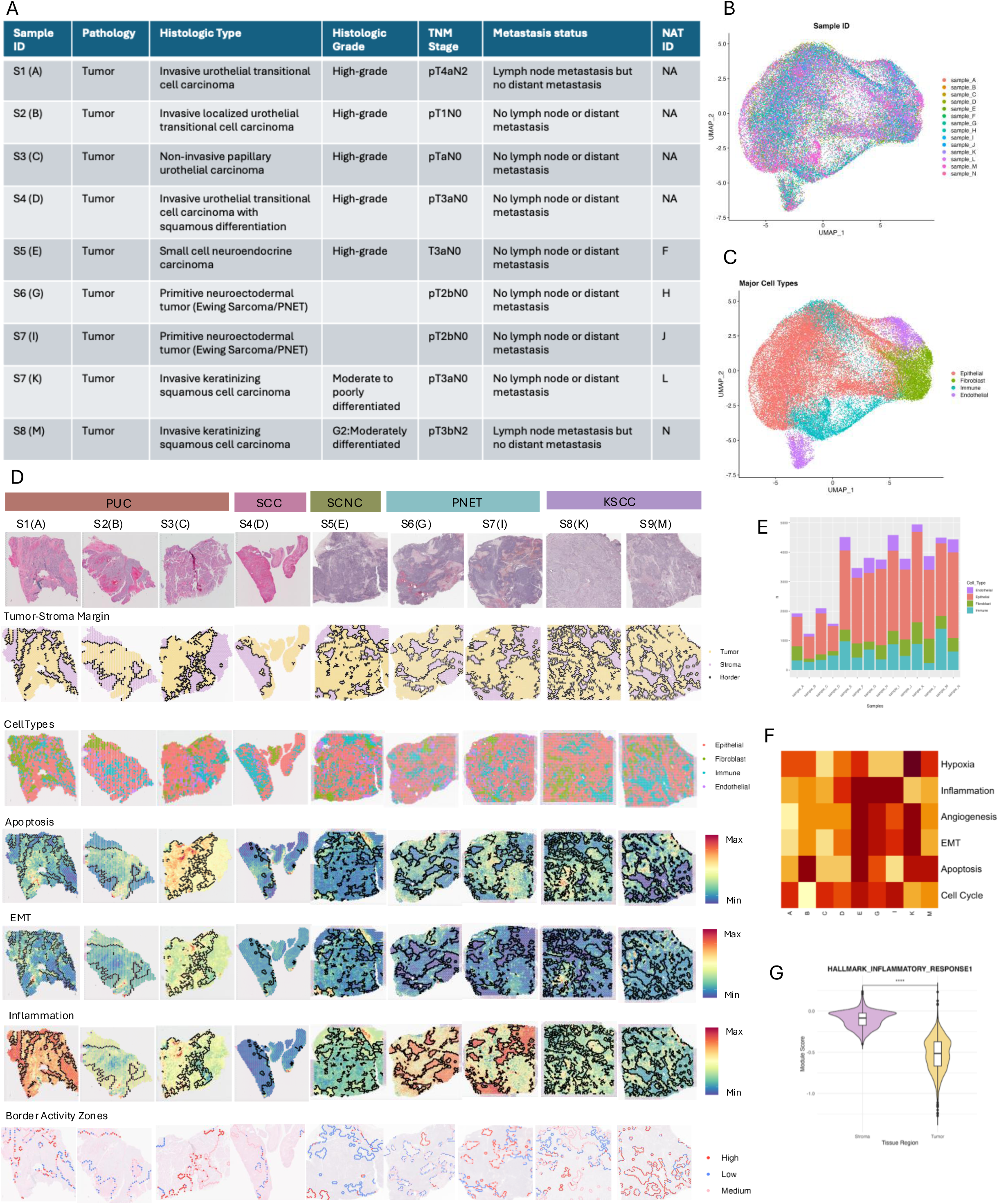
A) Table of sample information for in-house bladder cancer cohort (BC=9, Nat = 5). B) UMAP of integrated Seurat object for the BC cohort, colored by sample ID. C) UMAP of integrated Seurat object for the BC cohort, colored by major cell types. D) H&E staining images, tumor-stroma margins, major cell type plots, apoptosis, EMT and Inflammation signature pathways and border activity zones for BC cohort. E) Barplot of major cell types present within the cohort. F) Heatmap of autocorrelation (Moran’s Index) for cancer hallmark pathways, the darker the color the higher the index. G) Violin plot for pathway score for the inflammatory response pathway across stroma and tumor regions in sample A.

All samples showed different stages of aggressiveness that varied within and between subgroups. Our cohort consisted of mostly muscle invasive samples with only one non-invasive (Sample C) specimen. Tumors belonging to the rare histologies were of more advanced stage (all >T2b) when compared to the normal PUC. Sample A and M from the PUC and KSCC subgroups had the most advanced pathologic stage, T4 and T3 respectively, and with lymph node metastasis.

The histologies were initially determined during routine histopathological evaluation for clinical diagnosis on formalin-fixed, paraffin-embedded samples and were then reviewed by a pathologist. We annotated the nine samples based on H&E staining and their labels were later verified with the use of subtype specific cell markers during downstream analysis.

### 2.2 Heterogeneity in cell type composition within and between tumors

We performed SpaT analyses on a total of 9 tissue sections from 8 samples, that represented both common and rare histology of bladder carcinoma. One case had two different sections from the same primary tumor profiled and 5 samples had adjacent normal adjacent tissue (NAT) collected. After jointly processing the data from all samples in the cohort and correcting for any potential batch effect, no systematic technical variation was reported.

Following the standard Seurat workflow, UMAP dimensionality reduction was performed and used for identification and visualization of similar and different cell types across samples. Gene expression clustering of the spots identified 16 clusters (0-15). Coloring of the spots by sample ID verified the successful integration of all samples and showed no identifiable batch effect (**Figure 3B**).

Analysis of gene expression patterns of subtype specific gene markers was used for verification of subtype labeling. TP53 was used as control for the bladder cancer samples. As expected, all samples showed expression for this gene. The ESWR1 gene characteristic of the Ewing family of tumors showed no expression on PUC but was present in PNET subgroup. The KRT6A gene involved in the production of keratin, had high expression in the keratinizing tumors and little to no expression in every other sample. These observations helped us validate the labeling given through histopathological observations.

Module-A of CALISTA revealed a varied ratio of tumor to non-tumor regions across the samples. In PUC subtypes (A-D), a notable characteristic was the presence of contiguous tumor-rich regions. Conversely, rare tumor histology frequently displayed a more interspersed tumor-stroma arrangement, indicative of a marbled architecture. Sample A had the most stroma clusters from the PUC group. This was the only sample of PUC with lymph node metastasis. The more stroma invaded tumor phenotype could indicate a shift to a more aggressive phenotype.

Next, we annotated cell types and sub type lineages therein based on canonical markers and cluster-specific genes, and observed that epithelial cells, fibroblasts, immune and endothelial cells dominated the tissue microenvironments (**Figure 3C-D**). Different proportions of each cell type were present within each sample (**Figure 3E**), with most of the tumor being composed of epithelial cells, followed by immune infiltration. The heterogeneous compositions of the TME in bladder tumors across tumor tissues and normal tissues are consistent with a recent single-cell transcriptome study of rare bladder tumors [15]. Unsurprisingly, most epithelial cells lie within the tumor border with some sporadically spread throughout the stroma. Stroma sites were mostly composed of immune, endothelial and fibroblast sites clusters with some of these present inside the tumor zone. PUC sample B in particular had a high immune invasion inside the tumor.

CALISTA revealed a distinct distribution of active and passive zones within local tumor borders. KSCC sample M appeared to have the highest number of active borders. Like other samples, local margins were mostly surrounded by immune cells and fibroblasts. However, contrary to the rest of the cohort, sample M sample had minimum inflammation score in these stroma regions, where every other sample had its maximum inflammation scores at the immune and fibroblast clusters. It being the only sample among the rare subgroup with lymph node metastasis, this behavior might explain its aggressive phenotype.

On the other hand, samples B and D from the PUC cohort had the least number of active borders, which could be due to the high infiltration of immune cells within the tumor. Sample B contained a localized community of immune cells within the tumor core. The apoptosis score for this sample was significantly higher inside the tumor region when compared to outside the tumor and specifically concentrated around the immune cells. Inflammation scores were significantly lower inside this immune cluster when compared to other immune communities. Compared to other samples, the fibroblast cells aggregated at one single location at the northern most area of the slide. Margins surrounding this cluster had higher activity levels. The different distribution of stroma within this tumor supports the reduced amount of active interface regions.

### 2.3 Spatial heterogeneity in cancer hallmarks in tumors

Intra- and inter-tumor heterogeneity in cell type composition led us to ask whether the cellular processes and cancer hallmarks also varied within and between samples, especially considering the local tumor-stroma margins in a context-guided fashion. To assess this, we calculated the MSigDB pathway signatures for each spatial barcode in each sample and analyzed their distribution within the tissue. Here we discuss the pathway scores for cell cycle, apoptosis, hypoxia, inflammation, angiogenesis, and EMT.

We found the spatial distribution of the pathway signatures to show substantial spatial variation in the samples in our cohort. Pathway scores are calculated and scaled sample wise so, to compare between samples, we calculated the Moran Index for autocorrelation to evaluate the spatial clustering of these features and provide a quantitative measure of the captured aspects in our samples. A higher index indicates scores are highly similar between neighboring spots and that these features tend to cluster together within a tissue. Moran’s statistic for each pathway was calculated and represented in the form of a heatmap for easier visualization and comparison (**Figure 3F**).

First, we focused on the Cell cycle and Apoptosis score, since these are directly associated with tumor growth. We observed an inverse relationship between the autocorrelation scores. Samples having higher clusters of cell cycle activity had more dispersed apoptosis signals throughout the tissue. The exception being sample E which had a comparable score for both (Cell cycle: 0.54 and Apoptosis: 0.61). Analyzing the spatial location of this score revealed that both apoptosis and Cell Cycle scores were significantly increased inside the tumor region compared to its stroma counterpart. In particular, the apoptosis score was low and homogeneous across the stroma regions with clusters of high scores found within the tumor-rich regions (**Figure 3D**). The differences between the two regions are striking enough that tumor-stroma local borders almost perfectly segregate high and low scoring regions. The same could be observed at a smaller degree for the Cell Cycle scores, with mostly every sample having a significant difference between the two.

EMT and angiogenesis, both highly related to metastatic potential, were highly clustered (EMT: >0.68, Angiogenesis: >0.60) in most rare samples (E-K). Moran scores for these two pathways were very similar for each sample, with a difference of less than 0.05 in most cases. Identifying these clusters’ location inside the tissue, highly scoring niches of these features mirrored each other and were centralized within the tissue away from local borders (**Figure 3D**). The exception for this being sample E whose biggest cluster was found on the stroma region near highly active interfaces. Contrary to the big clusters of apoptosis and cell cycle homogeneously encompassing all the tumor tissue, EMT and angiogenesis niches were found in tight, well-defined clusters sporadically found inside the tumor regions.

Lastly, we analyzed inflammation and hypoxia which reflect the state of the TME. These two pathways appeared to be inversely correlated. High inflammation scores were found mostly on the stroma region, while hypoxia environments existed primarily inside the tumor tissue. Similar to the cell cycle and apoptosis score, the difference between the two regions was significant in almost every sample (**Figure 3G**). Calculated tumor borders perfectly segregate the slide into high and low scoring regions for these pathways. Basal samples (D, K, M) showed opposite behavior from the rest of the cohort, having low inflammation scores in stroma regions compared to tumor tissue. As expected, the autocorrelation for these was lower in this group, having a more random distribution of inflammatory niches. This compared to the high Moran’s I of other samples, particularly the neuronal samples, highlighting the inverse behavior of this subgroup.

Analysis of cancer specific pathways revealed distinct patterns across samples and histological subtypes. The diverse range within and among samples showed the high diversity present within a tumor. Only the Neuroendocrine sample (E) exhibited high clustering for all features (> 0.61), which implies a more heterogeneous environment when it comes to cancer functions involved throughout the tumor. The significant difference observed between tumor and stroma regions highlights the usefulness of cancer hallmark pathways when it comes to identifying tumor regions further validating the local margins identified by our method.

### 2.4 Histology-dependent patterns of spatial heterogeneity of tumor subclones

Tumors are clonally heterogeneous, where different clones may have significant differences in growth characteristics and interactions with the microenvironment, which in part, may influence the observed heterogeneity in cancer hallmarks at the boundaries and interiors of the tumors. Thus, we next examined clonal makeups in respective tumors. Setting the previously identified non-epithelial cells as reference, the copy number variation for each spot was calculated using the inferCNV software and the cancer clones were identified for each sample (**Figure 4A**). The dendrogram was cut to detect up to 5 major subclones for harmonized comparison between the samples. The tumor subclones had megabase-scale, or chromosome arm-level alterations, that distinguished them from the normal tissue, and from one another. The architecture of clonal phylogeny and proportion of spatial spots dominated by specific subclones in the tumor-rich regions varied across the samples and subtypes. As a negative control, running the inferCNV software on the NAT samples revealed no clonal lineages (distinguished by copy number alterations), verifying their labeling as tumor-free.

**Figure 4.**
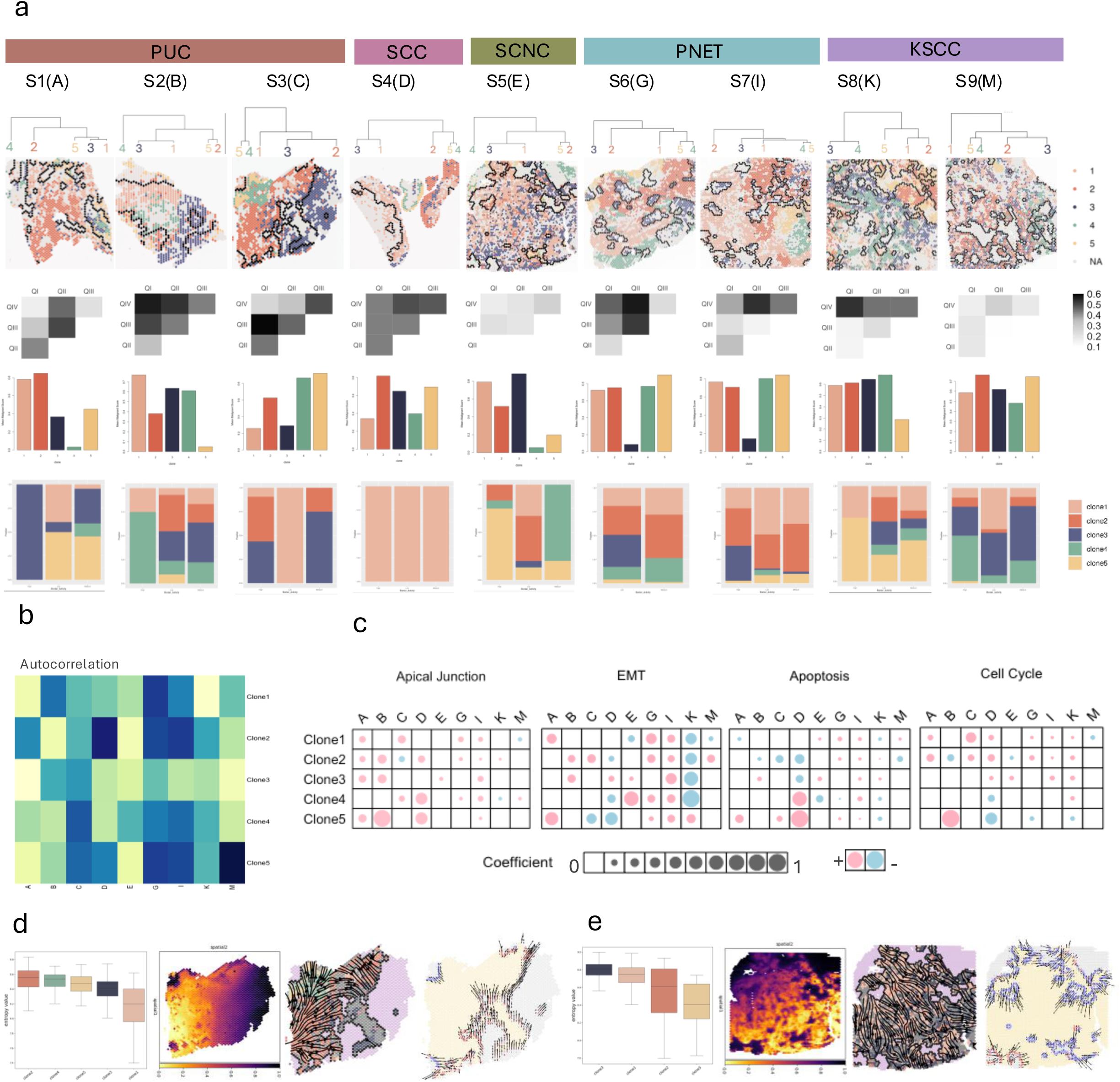
A) Dendrogram of cancer subclones, cancer subclones and tumor border spatial plots, heatmap for Bray–Curtis Dissimilarity score for subclones distribution across quadrants, barplots of mean malignancy score across subgroups, proportion of subclones present at the active, passive and intermediate zones. B) Heatmap for autocorrelation of clone subgroups per sample, the darker the color, the higher the Moran’s I. C) Bubble plot of significant regression coefficients for clonal subgroups and cancer hallmark pathways across samples. D) Entropy value of clonal subgroups, pseudotime plot of clone 2 and predicted lineage of clone 2 across the tumor tissue for sample C. Predicted trajectories of cells at the tumor-stroma border for sample C. E) Entropy value of clonal subgroups, pseudotime plot of clone 3 and predicted lineage of clone 3 across the tumor tissue for sample E. Predicted trajectories of cells at the tumor-stroma border for sample E.

To analyze spatial distribution of the subclones within respective tumors, the slides were divided into four quadrants and the Bray-Curtis Dissimilarity score for each quadrant was calculated. The results are represented as a heatmap, with the darker the color the more dissimilar those two quadrants are (**Figure 4A**). The overall Heterogeneity Score (HS) of the slide was calculated to be the sum of these, where a high value would indicate higher clonal heterogeneity. Certain patterns could be observed through the clonal distribution. As a general trend, the rarer histologies had a lower Heterogeneity Score (HS), with samples E and M having the lowest overall (E:1.17, M:1.11). Subclones in the PUC group on the other hand appeared to be more spatially segregated into their own groups, with sample D having the highest heterogeneity score (D: 3.34).

As observed previously, the rare histology exhibited a more marbled pattern of stroma niches within the tumor core while the PUC group was mostly composed of a continuous tumor mass bordering non-tumor regions. In most samples, most clonal subgroups were found at a higher concentration within the tumor core, with only one or two more at the non-tumor regions. As an example, see sample C, where almost every clone within the tumor border with only clone 3 outside. In most cases, then, clonal heterogeneity is also dependent on tumor to non-tumor distribution which varies within common and rare subgroups. These segregation between non-tumor and tumor regions in addition to the spatial divide among tumor clones influenced the final HS score.

Tumors with lymph node metastasis were more homogeneous in comparison to their none-metastatic counterparts from the same subgroup. Clonal distribution in sample M was more homogeneous than sample K (HS: 1.92). The same can be observed across the PUC samples where sample A has a difference of 0.5-1.3 in the HS when compared to the other samples within its group. This finding is consistent with the notion that in progressively advanced tumors, intra-tumor spatial heterogeneity decreases, at least locally, potentially owing to matrix remodeling and increased tumor cell mobility. Indeed, analyzing cancer cell-cell signaling, we found elevated matrix remodeling signaling in metastatic samples.

To quantify the level of spatial clustering among subclones, we once again calculated the Moran’s index for autocorrelation. Keratinizing and Neuroendocrine samples had a lower score on average. Samples with the lowest Heterogeneity score (E and M) had a low Moran’s I (<0.3) for most clones. On the other hand, clone groups in Neuroectodermal samples (G and I) had higher degree of clustering across all clonal communities, once again highlighting the inverse patterns observed between these subtypes. Since all of these are aggressive tumors, the compactness exhibited by the clone groups within the neuroectodermal samples suggests a divergent mechanism within this subgroup controlling the motility of its subclones and limiting their dispersion. While the opposite is observed for other samples, these signals indicate a structural difference across subtypes.

### 2.5 Subclones display divergent characteristics at local tumor-stroma interfaces and cores

We calculated the mean malignancy score for every clone group and noticed malignant signatures varied across clones (**Figure 4A**). Several samples showed a subgroup with significantly lower score compared to the others, closer to zero. Contrary to other groups, these subclones were found both inside and outside the tumor border, with the majority concentrating in stroma regions. We identified these clone groups as non-tumor normal epithelial cells. As an example, see sample A clone4 which has an almost zero malignancy score and is only present in the stroma region.

Next, we aimed to identify specific clonal groups within a sample whose behavior was different from other clones within the tumor. To identify these patterns, we performed spatial regression on the cancer hallmarks using each clonal community as the regressors. We decided to focus our attention on the apical junction, EMT, apoptosis and cell cycle pathway since these can give insight into the growth and dispersion of each group (**Figure 4C**). Not every clone had a significant coefficient for every pathway, clones which had been identified as normal epithelial cells or clonal groups with low Moran’s I had non-significant coefficients. This makes sense, since clonal groups with high random dispersion wouldn’t be identified as effectors of highly autocorrelated features and normal cells wouldn’t influence cancer signatures.

Regression analysis pinpointed clone 2 in sample C to be the only one with positive significant EMT coefficient (0.17). This led us to take a closer look into this clone and its interactions within the sample. SpaTrack [16] was used to calculate pseudotime, predict cell trajectories and evolutionary lineages of the clonal groups. Entropy analysis revealed clone 2 to have the highest probability for transition (**Figure 4D**). Predicted trajectories portrayed clone 2 as the starting clone within our tissue, with others clone lineages branching out. This clone also had a lower autocorrelation coefficient and its dispersion throughout the tumor mirrors the trajectory lines. Pseudotime plots showed stroma regions to be out of reach, further emphasizing the correct annotation of tissue regions. Clone 2 was found neighboring highly active boundaries (**Figure 4A**). Predicted transition direction at these regions shows border cells spreading towards the stroma (**Figure 4D**), further confirming its aggressive and invasive nature.

Previous studies had shown a correlation between high apical junction and EMT expression in relation to metastasis [17]. Clone 3 in sample E was the only clone with positive coefficient for both pathways within this sample (EMT: 0.02, Apical Junction: 0.01). Entropy analysis highlighted clone 3 as having the highest transition probability. Trajectory plots within the tumor show this group spreading all over the tissue. Additionally, this clone had a high positive coefficient for apoptosis and cell cycle portraying it as an aggressive clone with high turnover rate. While this clonal group was spread evenly throughout the tumor, there appeared to be high concentration in the top right corner near an active margin that will be examined at later sections.

The predicted direction for the transition of cells within the tumor border revealed opposite behaviors among passive and active zones. In sample C, passive borders appeared to be moving into the tumor while highly active regions moved into the stroma. The opposite was observed for sample E, where most cells within the active regions moved inward towards the tumor, the only notable exception being the right top corner surrounding clone 3 aggregates. This behavior led us to further classify active zones into progressive and regressive. Active regions in both samples contained clone groups with low malignancy scores, indicating the inverse behavior to be mostly attributed to cell types within the stroma. This motivated us to take a closer look at the minor cell types and the interactions occurring within the tumor stroma.

### 2.6 Differences in Minor cell types composition

We identified minor cell subtypes derived from the four major cell lineages using cell type and BC subgroup specific biomarkers (**Figure 5A**). Subgroup specific cells such as neuronal, neuroectodermal and differentiated keratinocyte were identified only on the corresponding tissue. On all other samples, we identified basal and luminal cells, cancer associated fibroblast (CAFS), such as Myofibroblast-like (Mcafs) and Inflammatory (Icafs) CAFS, and immune cells in the form of T cells, B cells and myeloid cells, specifically macrophages.

**Figure 5.**
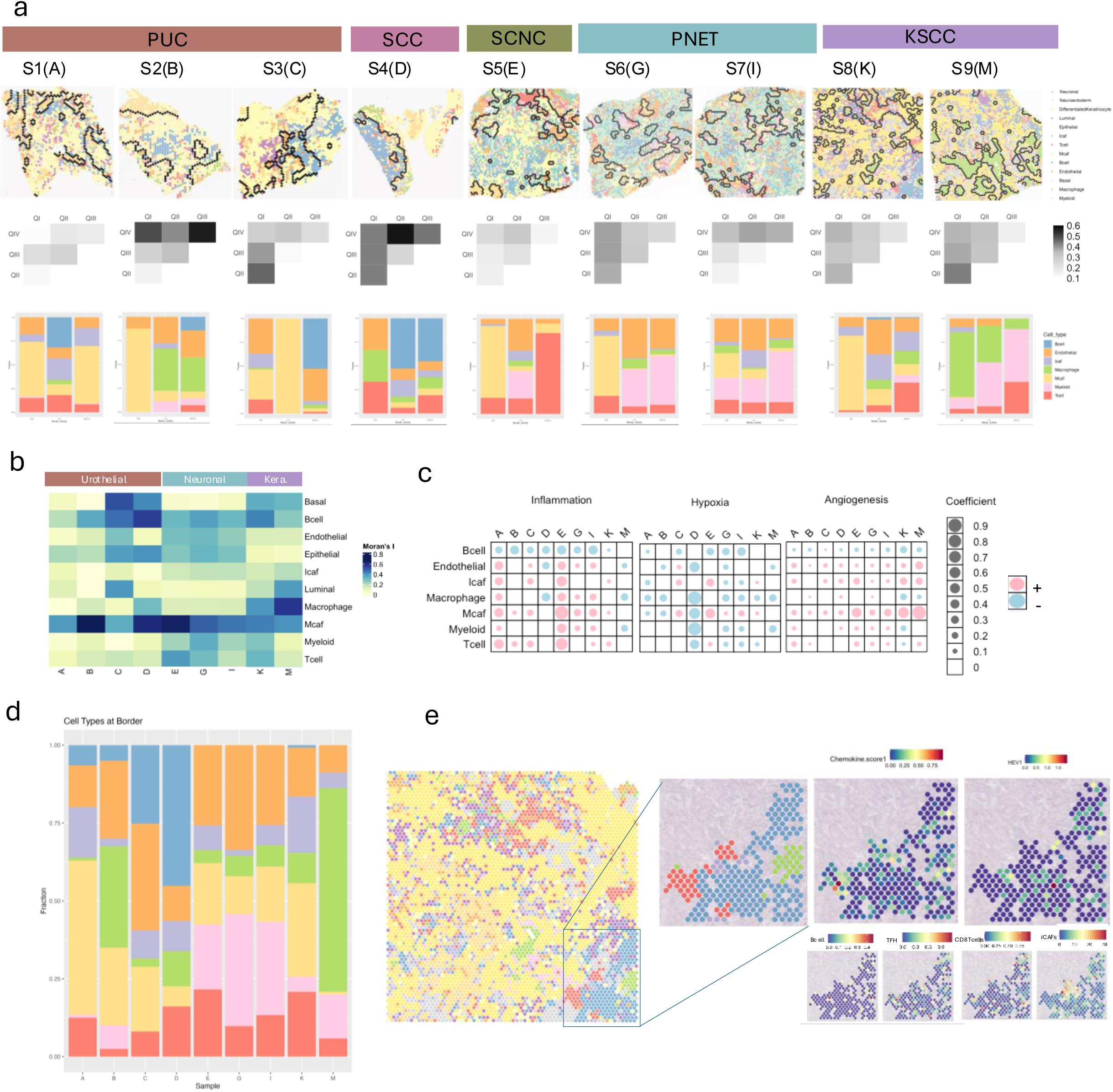
A) Spatial plots of minor cell types and tumor-stroma border, heatmap for Bray-Curtis Dissimilarity Score for cell type distribution across quadrants, barplots of minor cell type proportions across active, intermediate and passive zones. B) Heatmap of autocorrelation (Moran’s Index) value for minor cell types across the cohort. C) Bubble plot of significant regression coefficients for minor cell types and cancer hallmark pathways across samples. D) Stacked barplot of proportion of stroma cell types present at tumor-stroma border. E) Zoomed in of sample K showcasing TLS region. Plots showing chemokine, HEV, Bcell, Tfh cell, CD8 T cell, and iCAF signature score at TLS region.

Samples showed distinct cell type composition and grouping. Similar to previous analyses, we calculated the Beta diversity of every quadrant, the sample’s overall Heterogeneous Score, and the minor cell types autocorrelation score. Diversity patterns observed at the clonal levels were mirrored in the minor cell type composition. Once again, the rare subgroup exhibited higher homogeneity when compared to the common samples. In particular, non-metastatic PUC showed more spatial heterogeneity while the rarer and more advanced subtypes presented a more homogeneous mixture of the cell types.

Looking at the proportion of minor-cell types present within the tumor-stroma border, we noticed that active regions were mostly composed of Mcafs in most samples (**Figure 5A, D**). The major exception of this being samples D and M form the basal-squamous subgroup where M had mostly Macrophages and D a mixture of Macrophages and T cells. While other samples also had macrophages present at the tumor-border (**Figure 5D**), they were mostly present at passive or intermediate zones. Previously, we had highlighted how sample D differs from others in its low amount of highly active borders. Both passive and intermediate zones neighbor B cells, which explains the low cancerous activity observed at the local margins.

When analyzing the clustering of the minor cell lineages, we once again identified an inverse relationship between neuronal and keratinizing samples (**Figure 5B**). Cell types such as endothelial and epithelial cells had higher clustering in Neuronal subtypes while Basal, Luminal and macrophages showed higher clustering in the Keratinizing group. Mcafs and B cells showed a high degree of clustering across all samples. T cells were more spatially segregated in PUC compared to the rarer histologies. This, combined with the fact that rarer histologies contain a higher proportion of T cells and B cells implies higher immune invasion.

We repeated our spatial regression analyses, but this time focusing on different pathways (**Figure 5C**). For stroma cell type activity, we focused on the Inflammation, Hypoxia and Angiogenesis pathway which play a major role in the TME. We set minor cell types as the regressors and the respective pathway as the dependent variables. Macrophages are known to be key players in inflammation and as expected showed mostly positive coefficients for the inflammation score in most samples. Surprisingly, it had a negative coefficient on the basal-squamous samples M and D (−0.09 and −0.12 respectively). For basal-squamous sample K, the coefficient was also negative (−0.004), although not significant (p-value = 0.65). This combined with the fact that macrophages were highly clustered in these samples, but not on others, suggests an opposite behavior and role in the TME.

We observed T cells having mostly positive coefficients for inflammation across all samples. T cells, such as CD8+ T cells, secrete pro-inflammatory cytokines that attract other cells to sites of infection so it makes sense their presence would be a driving force in the overall inflammation score within the tumor. On the other hand, hypoxia coefficient results were overall negative. The only exception being Mcafs in the neuronal samples. Past studies [18] have shown that hypoxia drives fibroblast into their tumor-promoting CAF phenotype. The high coefficient of Mcafs for hypoxia in these samples could potentially indicate a higher CAF activity within these samples.

Finally, angiogenesis coefficient scores were overall positive and statistically significant in most samples for endothelial, Icaf and Mcaf cells. These three cell types have been widely studied and their pro-tumoral phenotype often includes the promotion of angiogenesis. Endothelial cells have a role in the formation of neovessels [19] and CAFs secretion of factors such as VEGF, PDGF and IL-6 promote this process [20]. Of particular interest though, is how sample E, K and M from the SCNC and KSCC subtypes exhibit a greater regression coefficient for CAFs compared to other samples. Having regression coefficients greater than 1, these three samples more than double the coefficient of the other groups whose value is well below 1. Sample M, with a coefficient of 4.63, exhibits a particularly high coefficient for Mcafs not seen in any other sample. This being one of the most aggressive samples with lymph node metastasis (T3bN2) suggests Mcafs to have a major role in its aggressive pro-metastatic signature.

### 2.7 Tertiary Lymphoid Structure Formation within Keratinizing samples K

When analyzing clustering patterns of immune cells within our specimens, we observed immune cells to be mostly clustered in the stroma and sporadically spread inside the tumor. Sample K however, contained a large cluster of immune cells inside the tumor core. This cluster contained both B cells and T cells with high autocorrelation scores (0.43 and 0.31 respectively) not seen in any other sample. This anomaly drove us to further inspect the immune cells and interactions occurring within this sample.

Upon further analysis, this cluster had features of widely studied immune organoids present in cancer known as Tertiary Lymphoid Structures (TLS). TLS structures have several stages of development, and the cell type composition and molecular signatures vary at each stage. We hypothesized this cluster may be an intermediate between an immature and mature structure based on its clustering and cell types. Early stages show a more dispersed mixture of both T cells and B cells, while more advanced stages contain packed groupings of these two.

Previous studies had shown that these organoid structures can be identified based on expression of 12 chemokines [21]. We calculated an aggregate score of these chemokines and were able to verify through expression that our structure was most likely a TLS (**Figure 5E**). Additionally, we gathered cell type-specific markers reported by other groups to be found within these structures [22]and calculated their aggregate score. Those included specific genes expressed by B cells, Tfh, CD8+ T cells and iCAFs inside TLS structures. Our sample showed expression for these genes. This also suggested the presence of a TLS structure in keratinizing sample K.

Studies have shown the presence of high endothelial venules (HEV) in TLS suggest an advanced stage of the organoid [23]. Using HEV specific markers identified by Sawada et al. [24], we identified the presence of these venules within our samples. The presence of TLS is closely associated with good prognosis among patients, so identifying this advanced structure inside this sample may indicate a better outcome, despite its aggressive phenotype.

### 2.8 Distinct Macrophage activity among BC samples

Previously, we identified that macrophages appeared to be highly clustered in the basal-squamous samples but with negative regression coefficients for inflammation scores. These two observations indicated different behavior of macrophages within this subgroup compared to the rest of the cohort. To verify this, we analyzed signaling interactions of macrophages and the cancer clone groups within these samples (**Figure 6A**).

**Figure 6.**
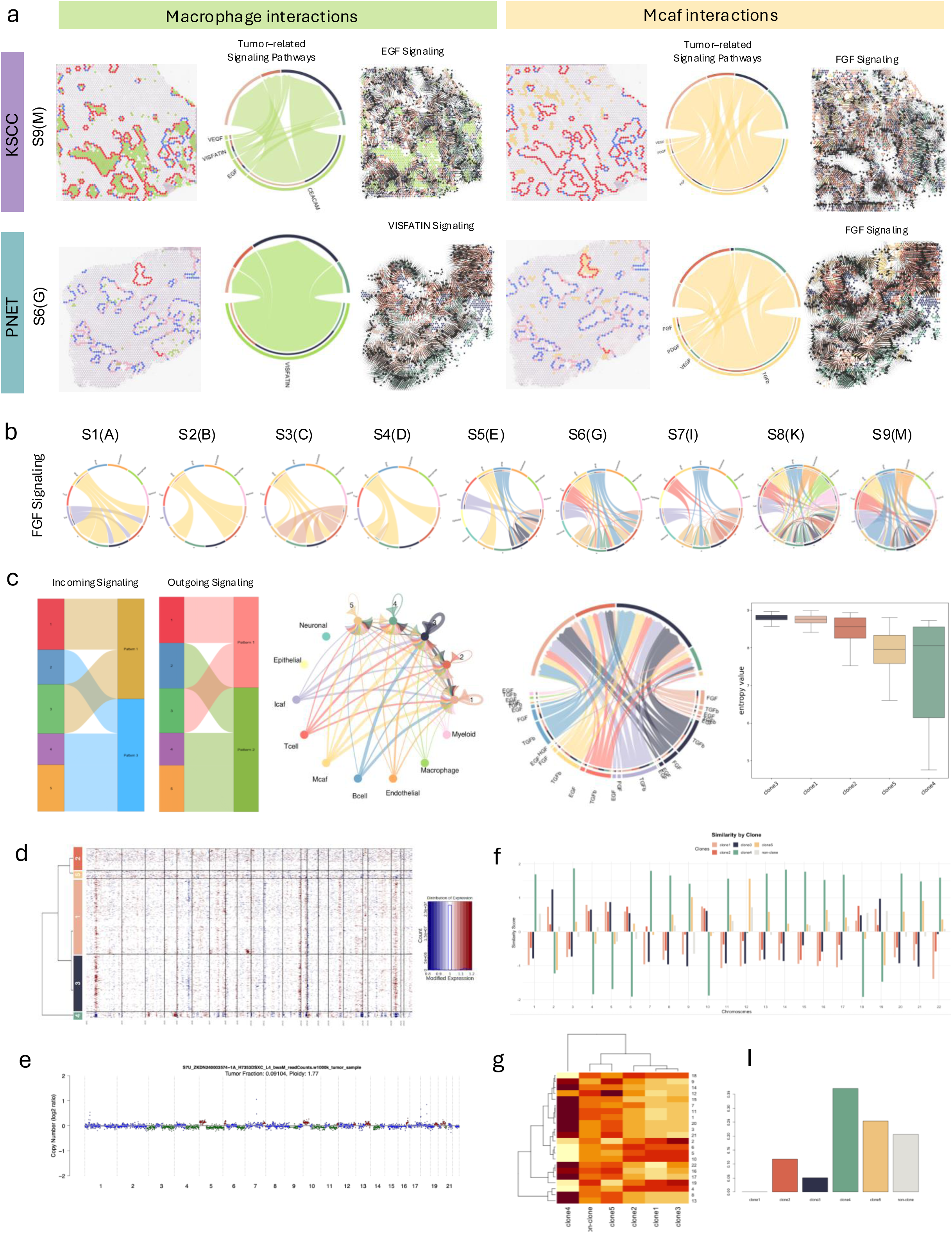
A) Comparison of Macrophage interaction and Mcaf interaction between sample M and G. Spatial plot showing spatial location in relation to border zones. Chord plots of signaling of pro-tumor pathways sent by macrophages and Mcafs towards cancer clones. Spatial plots showcasing direction of signaling for particular pathway and their spatial location within the samples. B) Circle plots showcasing signaling of FGF pathway. Sender being the stroma cell types and receivers being the cancer subclone groups. C) Incoming and Outgoing signaling patterns for cancer subclones in sample E. Circle plot showing total signaling received by cancer subclones within this sample. Chord plot showing incoming signaling from growth pathways towards cancer subclones. Entropy values of cancer clones in sample E. D) InferCNV copy number alteration plot from sample E. E) CNA plot from ctDNA data from sample E. F) Cosine similarity values by chromosome between expression patterns and CNA z-score. G) Heatmap of cosine similarity across clones in sample E. I) Barplot showing calculated relative contribution by clone to ctDNA liquid biopsy sample.

Studies have shown tumor associated macrophages (TAMs) have both pro-inflammatory Tumor Suppressor (M1) and anti-inflammatory Tumor Promoter (M2) states [7]. TAMs are highly plastic and transition between these roles continuously. Particularly in the context of BC, groups have highlighted how a high M1/M2 ratio is associated with favorable prognosis [25]. Here, we evaluated the presence of these two states among our samples to determine if this could be influencing tumor activity at the border. We selected known growth factor pathways (TGFb, VEGF, EGF) associated with M2’s tumor promoting activity and inflammation regulatory pathways (CEACAM, VISFATIN) regulated by M1’s signaling for our analysis.

Keratinizing sample M had many macrophages present in big clusters within the tumor stroma, mostly surrounded by active borders. Signaling analysis revealed high interaction between the macrophages and clones 1,2,3 and 4. Clone group 5 didn’t have any statistically significant pathways which made sense given that cells from this clone group are clustered farther away from the macrophage communities. We took a closer look at the EGF pathways since studies have shown signaling from macrophages to the EGF receptor increases tumor growth and metastatic potential. Spatial analysis of the EGF signaling pathway revealed a high amount of signaling with clones 2,3 and 4 which mostly compose the active border zones (**Figure 4A**). The increased interaction of EGF and other growth factor pathways could potentially explain the high activity seen within these margins.

Samples outside the KSCC subgroup didn’t contain macrophage clusters at the tumor-stroma. Sample G presented with more dispersed distribution of macrophages which didn’t appear to have a significant impact on border activity. From the pathways used in our signaling analysis, only the VISFATIN pathway exhibited any signaling towards cancer clones within the sample. The clone groups and regions receiving these signaling correlated to regions with high inflammation previously shown (**Figure 1**). These observations highlight Macrophages to have more of an M1 pro-inflammatory phenotype in this sample compared to the M2 tumor promoting activity previously seen in sample M. Similar behavior in other samples, with only samples in KSCC showing activity in tumor promoting pathways.

### 2.11 Tumor Associated Fibroblast promotes tumor activity at Tumor-Stroma Border

While macrophages were mostly dispersed in the PNET sample G, myofibroblasts (Mcafs) were highly clustered in the only active zone among the local tumor borders identified within this sample. Previously, we observed how CALISTA revealed Mcafs to be present at higher proportions in active border zones for almost every sample. Studies have shown Mcafs to have tumor-promoting activity in various cancers including BC [26, 27]. To verify whether this was the case within our samples we repeated our signaling interaction analysis previously performed on macrophages but instead selected Mcafs as the senders and the cancer subgroups as the receivers. We focused on tumor promoting pathways known to be upregulated by Mcafs such as FGF, TGFb, VEGF, PDGF and HGF [28].

Mcafs in sample G had significant interaction for 4 out of 5 of the analyzed pathways with most clone groups. Clone group 3 had the least amount of signaling. This being the group composed of mostly normal epithelial cells it was not surprising to see interactions toward this group to be decreased. Samples from various subgroups (A, B, K, E) also contained regions of high activity at the tumor - stroma border composed of Mcafs and significant signaling from many of the Mcaf associated pathways with almost every clone group inside the tissue. Only sample M didn’t contain any non-tumor regions mostly composed of Mcafs, as these were clustered primarily within the tumor core. However, the Mcafs present inside the tumor regions also exhibit similar signaling patterns to the other samples, particularly targeting clones 1 and 4. This, in conjunction with the signaling received by the M2 macrophages, had cancer clones receiving growth factors from both ends, which could further explain the aggressive phenotype we’ve observed throughout this study.

The FGF pathway was of particular interest to us, since studies have shown FGFR inhibitors are approved for the treatment of BC [29]. The FGF pathway had significant signaling in all our samples (**Figure 6B**), often active in most if not all of the cancer clones within the tissue. While less aggressive samples (B and C) had signaling coming only from Mcafs, more aggressive samples had other cell types contributing to the overall signal received by the cancer clones. In particular, highly aggressive samples K and M (T3) from the keratinizing subgroup have almost every cell type contributing to this pathway. This emphasizes the role of Mcafs and its effects on the TME and suggests the potential for BC patients to undergo FGFR inhibition therapy.

### 2.13 Identification of dominant subclones through cell signaling

From the previous signaling analysis, we noticed that signaling coming from within the TME was not consistent across all clones. The circle plots for sample E revealed clones 1, 2, and 3 with the highest amount of growth factors in the FGF pathway (**Figure 6B**). Sankey plots from the overall signaling patterns of the clonal communities within this sample clustered clone 1 and 3 into receiving a similar signaling pattern. This pattern contained several cell growth pathways (EGF and HGF) that were not present in the patterns observed by the other clones within this sample.

These two clonal groups also exhibited the highest malignancy score and had been grouped into the same branch in the dendrogram obtained during the initial CNV analysis (**Figure 3**). Circle plots showed the highest amount of interaction coming from any other cells in the TME and the highest amount of signaling coming from all growth pathways analyzed (FGF, TGFb, EGF, HGF). Entropy analysis also revealed these two clones to have the highest entropy within this sample, suggesting a higher potential from mobility and migration. All these observations suggested these two cancer clones to be particularly aggressive and dominant within the tissue.

### 2.13 Subclone-specific copy number signatures in urine samples

In addition to the tumor biopsy samples, we collected a urine sample from patient E. We hoped to identify the cancer signatures observed within the solid tissue using the liquid biopsy. Copy Number Alterations (CNA) were calculated from the cfDNA extracted from the urine (**Figure 6E**). Alterations observed in the cfDNA highly mirrored altered expression patterns observed in clone group 3 and 1 used to predict the copy number variations when first identifying the subclones through the InferCNV package (**Figure 6D**). Both datasets were analyzed in a chromosome-wise manner and the similarity between solid tissue expression and cfDNA CNA were calculated.

This analysis revealed clone 4 to have the most similarity to cfDNA in most of the chromosomes (**Figure 6F**). Clone 4 is the clone group within this sample mostly composed of normal non-cancerous cells. Additionally, the chromosomes in which this group presented the highest similarity were those with the least CNA. Hierarchical clustering identified subclones 1 and 3 to have comparable similarity scores (**Figure 6G**). Both had high similarity scores in chromosomes which presented a greater number of alterations in the CNA plot. It is worth noting that clone 2 which followed subclones 1 and 3 in both growth signaling and malignancy scores also had high similarity scores in these chromosomes.

When calculating the overall relative contribution of cancer clones to the liquid biopsy data, we found clone groups composed mostly of normal cells to be predominantly contributing to the signals found within the liquid biopsy sample. However, the cancer clones 1, 2 and 3 were still observed, even at a lesser degree and their presence confirmed through the similarity scores at genomic regions with high variation. Comparison of clonal expression to cfDNA’s CNA supports the identification of clone 1 and 3 as dominant and aggressive subclones within this tissue. Additionally, the identification of clonal signatures from solid tissue in liquid biopsy opens an avenue for monitoring and tracking tumor activities in a noninvasive way.

## 3. Discussion

In this study, we provided a comprehensive spatial transcriptomic characterization of common and rare bladder cancer histologies. Using our novel computational genomics framework, CALISTA, we reliably identified tumor borders and mapped the cellular interactions occurring at the tumor-stroma-immune interfaces. We demonstrated substantial intratumoral and intertumoral heterogeneity across BC hitologies, detailing the complexity of bladder cancer biology beyond the conventional histologic classification. By classifying tumor border as active or passive we identified clinically relevant TME niches composed of aggressive tumor clones, CAFs, and TAMs, which collectively amplify cancer signaling at the interfaces and promote tumor invasiveness. Importantly, several of the signatures identified at these tumor-stroma interfaces were also detectable through non-invasive liquid biopsy of urine. This highlights the potential opportunity to monitor these tumor border dynamics over time. Furthermore, our findings revealed substantial histologic-specific differences in TME cell type composition, spatial heterogeneity, signaling and tumor dynamics, suggesting divergent mechanisms of microenvironment remodeling that underlies the biological diversity across bladder cancer histologies. Collectively, these results stress the need for integrative spatial and molecular analyses to identify therapeutic susceptibilities within the ecosystem of the tumor-stroma-immune interface.

Through our analyses, we demonstrated that elevated tumor-promoting interactions between cancer subclones and non-tumor cells are enriched at active tumor borders. Subclone-specific upregulation of key ligand-receptor pairs revealed distinct roles for TAMs and mCAFs across the BC samples. We observed that mCAFs consistently formed dense clusters within stromal regions adjacent to active tumor borders, regardless of histology. We also found increased malignant signatures in subclones proximal to these aggregates. Consistent with prior studies, we identified the FGF pathway as a prominent signaling pathway mediated by CAFs, whose upregulation leads to tumor progression [30]. In our cohort, FGFR interactions were upregulated across histologic types. This findings has important implications, as FGF targeted therapies are already approved for metastatic urothelial carcinoma and are being investigated in earlier disease stages, suggesting potential applicability to variant histologies in addition to PUC [31, 32].

In contrast, TAMs showed diverse behavior across BC histologies. Notably, squamous cell carcinoma samples had larger TAM clusters localized outside the tumor borders, with evidence of increased M2 polarization. TAMs have gained growing attention for their tumor-promoting roles, including suppression of the anti-tumor immune response and facilitation of tumor progression. Prior studies have identified the squamous histology to be particularly aggressive, associated with poorer prognosis and reduced responsiveness to immunotherapy [33–35]. Together with our findings, this suggests TAMs play a unique role contributing to the aggressive phenotype of SCC and represent a potential immunotherapeutic target, in-line with the burgeoning field investigating TAM-reprogramming agents in oncology [36]. When analyzing TAM signatures in our paired PNET samples, we observed similar signatures within the tissues, further validating histologic specific signatures. Our analysis indicates that variant histologies may benefit from histology-specific therapies, and that spatially resolved analyses such as ours can inform a more personalized treatment approach.

Using CALISTA, we comprehensively characterized and compared both rare and common bladder cancer histologies to reveal distinct and shared features of their TMEs. These spatial and transcriptomic differences provide insights into the cellular dynamics that may influence the biological behavior and clinical management of each histologic variant. The samples analyzed in this study differ both in the cell of origin and by the phenotypic expression. Neuroendocrine carcinoma can arise from adenocarcinomas through transdifferentiation of tumor cells into neuroendocrine phenotypes, or from pre-existing neuroendocrine cells within the bladder epithelium [37]. Primitive neuroectodermal tumor of the bladder belongs to the Ewing family of tumors, typically with a fusion gene involving EWSR1 and the ETS gene family [38, 39]. These cancers originate from neuroectodermal precursor cells within the bladder’s nervous system network. Keratinizing squamous cell carcinoma in contrast, arises from SCC and is characterized by production of keratin, a fibrous structural protein normally found in skin, hair, and nails [40]. The diagnosis of these histologies, either in isolation or with de-differentiated UC, generally confers a poor prognosis and necessitates therapeutic escalation or more invasive interventions compared to those typically recommended for patients with PUC. Our analysis demonstrates that beyond the biomolecular differences, these histologies exhibited significant differences in cellular composition, spatial architecture and tumor-stromal-immune cell interactions. By analyzing this broad repertoire of bladder cancers histologies, we gain a more comprehensive understanding of the mechanisms underlying disease heterogeneity and progression across histologic variants.

This was a limited cohort, reflecting the rarity of these histological variants. Nevertheless, the trends observed within and across samples provide valuable insights into the mechanisms involved in the progression of these tumors. Even within PUC, and across different clinical stages of the same histology, comprehensive analyses have shown significant molecular heterogeneity that may explain stage-specific differences in risk classification [41, 42]. However, the complexity of the TME and its dynamic interactions remains incompletely understood. SpaT offers a window into the geographical composition of a neoplasm at a specific point in time. As shown by our results, these carcinomas are profoundly heterogeneous, and any given spatial snapshot may not completely capture the temporal evolution or overall composition of the tumor. While the use of liquid biopsy, specifically circulating and urinary tumor DNA has gained traction in bladder cancer as a marker of residual disease [43, 44], coupling such liquid biopsy analyses with spatial frameworks, like CALISTA, may yield deeper and more granular insight into tumor biology and microenvironment remodeling that cannot be captured solely through SpaT analyses alone. Additionally, we recognize how the incorporation of additional ‘-omics’ data can help further characterize the microenvironment of these tumors, and we look forward to integrating them in future studies.

In this study, we showcase the utility of CALISTA by applying it to bladder cancer histological variants. However, the method is cancer type agnostic and can be applied to analyze other malignant neoplasms and can utilize other spatial transcriptomic platforms. Our TLS analyses suggest that the resources could be repurposed to examine other localized tissue-level immune and other functional elements, perhaps in non-malignant conditions as well as development and aging.

## 4. Methods

### 4.1 Sample Collection and Acquisition

Deidentified human bladder cancer tumor tissue samples were collected with written informed consent and ethics approval by the Rutgers Cancer Institute of New Jersey Institutional Review Board under protocol no. Pro2019002924 (PI: De). We performed SpaT analyses on a total of 13 slides in a set of two batches. The first batch consisted of the common bladder cancer histologies as previously published [45]. Briefly, 5 μm tissue sections were placed on the Visium Spatial Gene Expression Slide for FFPE following Visium Spatial Gene Expression for FFPE-Tissue Preparation Guide (10X Genomics, CG000408)[46]. Each slide had 4 capture areas and each capture area was 6.5 x 6.5 mm with ∼5000 spots per capture area. Each spot was 55 µm in diameter with a 100 µm center to center distance between spots. Slides containing the tissue sections were incubated at 42 °C for 3 h and dried overnight at room temperature. Deparaffinization was then performed following Visium Spatial for FFPE – Deparaffinization, H&E Staining, Imaging & Decrosslinking Protocol (10X Genomics, CG000409)[47]. Slides were then used with Visium Spatial Gene Expression for FFPE User Guide (10X Genomics, CG000407)[48] to generate Visium Spatial Gene Expression – FFPE libraries and sequenced on Illumina NovaSeq S4 300 cycle. Outhouse pancancer samples were downloaded from 10xGenomics [49–51] and annotations of H&E images retrieved from available studies [8].

### 4.2 Seurat Preprocessing

The sequence data (FASTQ files) were processed using Space Ranger (v2.0.1) [52] count pipeline for single-library analysis of fresh frozen (FF) and formalin-fixed paraffin embedded (FFPE) samples to align transcriptomic reads to the human reference genome (GRCh38), map them to the microscopic image of the tissue from which the reads were obtained and generate Feature Barcode matrices. The resulting count matrices and associated H&E physiological images were then used by the R package Seurat (v4.3.0) [11, 53] for standard spatial transcriptomic analysis using default Seurat parameters; barcodes with less than 300 genes were excluded. Seurat v3 [54] was used for sample normalisation (SCTransform, mitochondrial and ribosomal mapping percentage were regressed out). In the first batch [45], we noted medians of 4877 genes per spot, and 12416 UMIs per spot. The medians for this second batch were of 5009 genes per spot, and 10743 UMIs per spot. All samples were uploaded as individual Seurat objects and integrated into a single object for easier analysis and comparison across samples.

The BC datasets were integrated using the Seurat SCTransform integration workflow (anchor-based method with 3000 variable genes), followed by principal component analysis (PCA, using first 30 principal components), UMAP dimensionality reduction (UMAP) and clustering. Dimension reduction and clustering analysis revealed no batch effect.

### 4.3 Active Tumor Border Identification

The R package SpaCET() [8] was used to determine the overall malignancy score from Visium samples. The SpaCET.deconvolution() function was used with cancerType parameter ‘BRCA’ for inhouse common subtypes, ‘PANCAN’ for inhouse rare subtypes and ‘BLCA’, ‘GBM’ and ‘PAAD’ for the outhouse samples respectively. The sfdep() package was used to calculate the Getis-Ord* [12] statistic on the malignancy score. Spot coordinates were retrieved from the Seurat object and the neighborhood object was created using sp()[55, 56] and spdep() [57] packages. Neighbors were defined as spots which lie in a radius of the minimum distance between all spots across a particular tissue. On average, each spot had four neighbors. The Gi* score was calculated using the local_g_perm() and ran on 999 simulations. Spots with significant positive scores were considered tumor rich regions, while those with significant negative scores were considered tumor poor regions. Spots whose p-value was not significant were labeled as interface.

To classify the interface region into the stroma and tumor compartment, we tested 9 different models. Each model consisted of a neural network classifier with 3 hidden layers, had tanh activation functions and was trained on 1000 epochs. The classifier was created using the h2o.deeplearning() function from the h2o()[58] package in R and tested with 9 different inputs. Each model varied in input features consisting of a mixture of spot coordinates, malignancy score, Cancer Hallmarks Pathways Scores and Differentially Expressed (DE) genes between the tumor-rich and tumor-poor compartment. The MsigDB [59, 60]Cancer Hallmark Signature gene sets were retrieved using the msigdbr() package and pathway scores calculated running the AddModuleScore() on the Seurat object. Cancer Hallmark scores, spot coordinates and malignant scores were used as input for the classifier. DE genes were calculated using the FindMarkers() function of Seurat with cell identity 1 and 2 being ‘Tumor-rich’ and ‘Tumor-poor’ respectively. The results were filtered to keep only genes with avg_log2FC > 2 or avg_log2FC < −2 and with p_val_adj <= 0.01.

The identified tumor rich and tumor poor spots were used to train the model on a 80:20 train and validation split. Model performance and variable features were extracted using h2o’s performance() and varimp_plot() functions. The model assigned interface spots to stroma or tumor regions. The tumor-stroma margin was defined as spots with both tumor and stroma neighbors.

Continuous margin borders surrounding stroma or tumor islands were identified as regions of interest using sp and spdep neighborhoods and adjacency matrices. Cancer Hallmark scores from these regions were extracted from the object and used as input for a clustering analysis. K-Means clustering was performed using the flexclust () [61] package with parameter k = 3. Scores in aggressive cancer activity pathways, such as EMT and Angiogenesis, were aggregated and clusters were labeled to High, Medium and Low based on the activity score

Differentially expressed genes between High and Low active interfaces were retrieved using FindMarkers function in the Seurat package. Genes with a significant p value and a log fold greater than 0.58 were extracted for gene ontology analysis. The enrichGO function of the DOSE [62] package was used to identify upregulated pathways. The presence of malignant and aggressive pathways in the GO analysis indicated successful identification of High and Lowly Active interfaces.

### 4.4 Cell Type Assignment

Major cell types were assigned to each UMAP cluster on the integrated object. We utilized known cell type specific markers from past studies and available databases for cell type annotation. Cluster specific marker genes were used for validation of correct assignment. Minor cell types shared across all samples were inferred on the integrated object. Subtype specific cell types were determined in a sample-by-sample manner. We used Spotlight’s [63] deconvolution to verify cell type assignment.

### 4.5 Cancer Subclone Identification

The R package inferCNV [64] was used to identify the subclonal communities. The Immune, Endothelial and Fibroblast population was chosen as reference cells. Hierarchical clustering was performed based on the transcriptomic profile and each dendrogram was cut at 5 branches to create 5 clonal subgroups for each sample. We arbitrarily chose an equal number of clone subgroups in each sample for easier analysis and comparison between the samples.

### 4.6 Spatial and Diversity Statistics

We represented the spatial map of the tissues with a neighborhood graph using the packages and method explained in section *4.3* *Active Tumor Border Identification.* Autocorrelation of cell-types and pathways was performed on the generated graph structure using the moran.randtest() function from the adespatial [65] package. We calculated the spatial regression of cell types with respect to cancer pathways using a spatial lag model (lagsarlm), as implemented in the spatialreg R package [66], to perform multivariate regression using a published approach. When performing regression using the clones as the regressors, we included inflammation and hypoxia as additional regressors to account for TME behavior. The epithelial score obtained through ModuleScore() was multiplied to the binary identity of each clonal community to account for other non-epithelial cells within a spot. When regressing on minor cell types, only the cell type prediction score was used.

We analyzed the spatial heterogeneity of clones and minor cell types by computing the beta diversity statistic of every quadrant of the slide.

### 4.7 Cell-to-cell communication analyses

The R package CellChat (v1.5.0) [9]was employed to analyze cell-to-cell communication between tumor cells and other cell types. We used the CellChat human database without additional supplementation. Preprocessing steps were all conducted with default parameters. The functions computeCommunProb and computeCommunProbPathway were applied to infer the network of each ligand‒receptor pair and each signaling pathway separately. A hierarchy plot, circle plot and heatmap were used as different visualization forms. For Spatial visualization of cell-to-cell communication, the python package COMMOT [67] was utilized using the same database and default parameters.

### 4.8 Clonal Trajectories

The python package SpaTrack [16] was used to determine the trajectories of the clones and borders in our samples. The starting cluster was chosen as that with the highest entropy value. The RNA velocity and pseudotime were calculated using spaTrack.get_ptime() and spaTrack.get_velocity() functions with default parameters. The entropy value and associated plots were also obtained through this package. For clonal trajectories, the starting clone was chosen based on its entropy value and validated through inferCNV dendrogram. The trajectories were drawn only on the tumor region. For tumor-stroma border trajectories, the tumor region was chosen as the starting cluster. Arrows embeddings are only shown for the tumor-stroma borders.

### 4.9 Clonal contribution to cfDNA

We collected paired urine samples from the patients, when available. Cell-free DNA was isolated from 2ml urine samples using QIAamp Circulating Nucleic Acid Kit, following manufacturer’s protocol. Low pass whole genome (3-5X) sequencing of cfDNA was performed on the Illumina platform after DNA-level quality checks. 150bp paired end reads were mapped to the human genome (hg38). IchorCNA (v0.3.2) [68], a probabilistic model designed for low pass WGS data from cfDNA, was used to identify copy number alterations at a megabase scale resolution and accordingly estimate tumor fraction and ploidy. IchorCNA employs a hidden Markov model (HMM) framework that fits observed read-depth signals to mixtures of tumor and normal components, accounting for GC-content and mappability biases. Analysis was performed using default parameters, briefly, bin size = 1 Mb, GC correction enabled, and ploidy range initialized between 1.6 and 3.0. The algorithm iteratively optimized likelihoods to estimate the most probable tumor fraction and segmentation breakpoints. Genome-wide CNA plots were generated using the IchorCNA_plot.R script and the ggplot2 R package (v3.4.0)[69]. CNV segmentation and genome-wide log₂ ratio profiles were manually reviewed to ensure smoothness, accurate breakpoint detection, and the absence of artifacts arising from low-mappability or low-coverage regions before integration with transcriptomic data. These metrics were used to assess the quality and interpretability of cfDNA-derived CNV profiles.

For sample E, which had paired spatial transcriptomic and cfDNA sequencing data available, we compared the genome-wide cfDNA copy number profile with that for subclones identified in the tumor tissues, as indicated above, using cosine similarity. The subclones showing relatively higher cosine similarity were highlighted.

## Acknowledgement

The authors acknowledge scholarly discussion from other members of Rutgers Cancer Institute.

## Competing interest statement

The authors have no competing interest.

## Author contribution

L.Q.Z. designed experiments, performed analyses, interpreted the results, and wrote the manuscript with input from all authors. S.B. performed clinical analyses and interpreted the results. A.B. processed biospecimen and performed genomic analysis. V.P. provided assistance with clinical sample collection, and interpretation of results. G.R. provided pathological annotation and interpreted the results. S.G. coordinated clinical sample collection, interpretation of results. S.D. conceived the idea, designed experiments, interpreted the results, and wrote the manuscript with input from all authors.

---

**Workflow.**
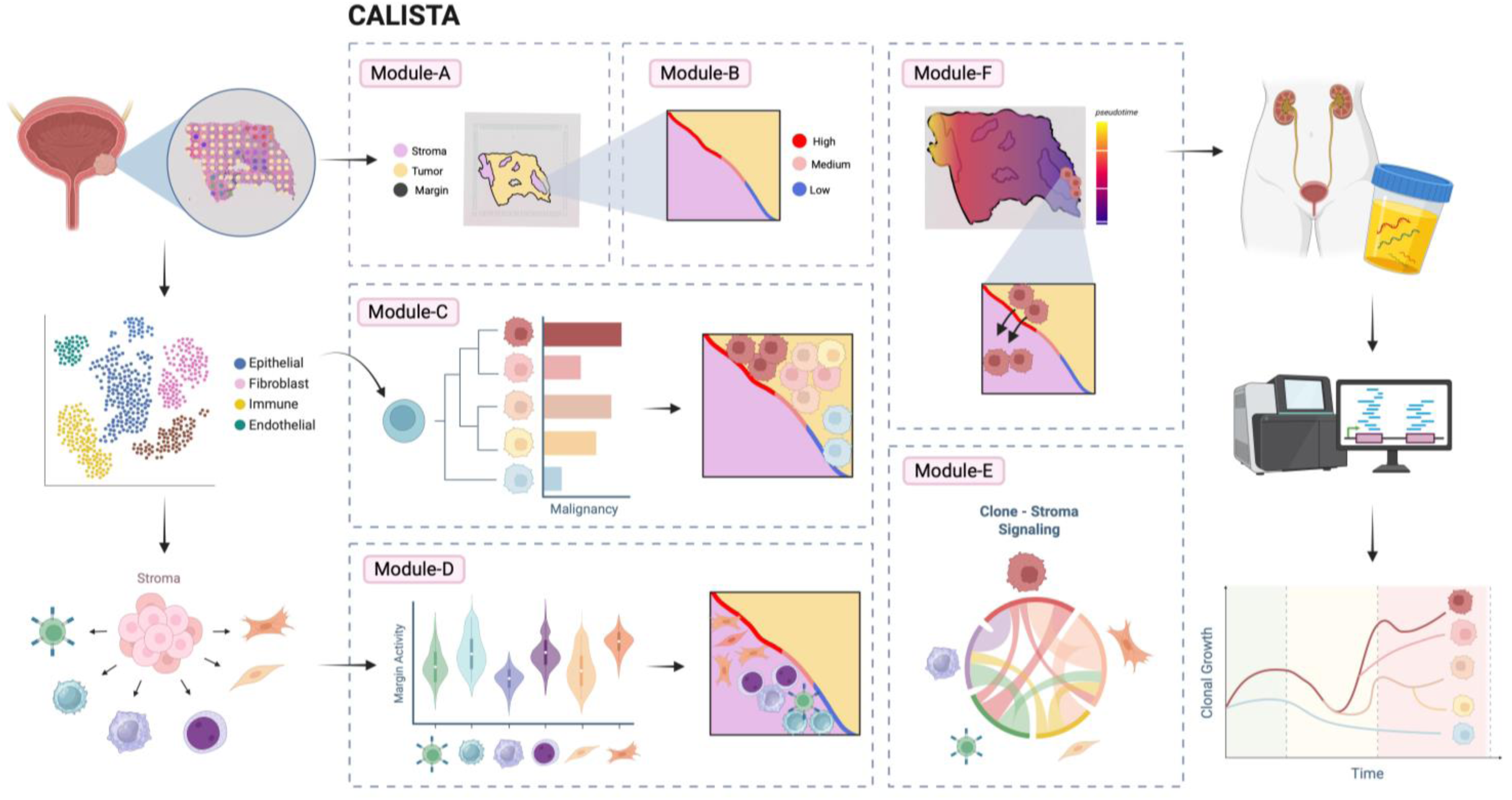
CALISTA has six main modules. ModuleA, identification of tumor-stroma borders. Module B, classification of tumor borders into active, passive and intermediate zones. Module C, inference of cancer clones and their localization within the tumor border. Module D, identification of stroma agents and their localization with respect to tumor border zones. Module E, clone-stroma signaling interaction analysis across stroma-tumor borders. Module F, pseudotime analysis of clonal evolution and identification of aggressive clones with potential for further invasion.

## Notes

### Competing Interest Statement

The authors have declared no competing interest.

